# Sample-Based Training Data for Effective 3D Cell Segmentation

**DOI:** 10.64898/2025.12.05.692330

**Authors:** Adam Smith, Till Bretschneider

## Abstract

Deep learning remains the leading choice for cell segmentation, a critical step in bioimaging analysis. Whilst deep learning models provide excellent segmentation accuracy, a large number of hand-annotated samples are required which are scarce in 3D. We propose a method for generating synthetic data for training deep learning models to augment or replace such datasets. By preserving key features in real data, we show that a standard UNet trained on synthetic data can segment single motile cells with branching filopodia with high accuracy. We also demonstrate how low-effort slice annotations can sufficiently replace volume annotations in our data generation pipeline. Overall, we provide a simple alternative to annotating large 3D datasets for training neural networks to segment cell imaging data.

## 1 Introduction

The demand for novel analytical techniques has grown in recent years due to the availability of large-scale 3D bio-imaging datasets of cellular and tissue-level processes. Accurate cell segmentation is a critical stage of analysis, but can be challenging due to the size, complexity and heterogeneity of the data. Several studies have highlighted that deep learning outperforms traditional methods for segmenting datasets with such variability, but require a significant amount of labelled training data [1, 2].

While hand annotations are the current gold standard, a recent study on breast cancer cells highlighted that hand annotations vary between individuals depending on their expertise [3]. Hence, majority voting schemes are often used to circumnavigate annotation variability across individuals. Since hand annotation is particularly time consuming for 3D imaging, majority voting is not often feasible in those studies.

To navigate the challenges in acquiring accurately labelled imaging datasets, several studies have proposed the use of synthetic alternatives. Such data can be produced by generative [4] or computational models [5, 6, 7]. Synthetic datasets offer distinct advantages for training segmentation models since they enable the generation of large datasets required for deep learning, allow controlled variation of image features to encompass real-world cases, and can be tailored to ensure balanced representation of rare classes. The method involves generating images from reference labels, avoiding subjective hand annotations which are known to adversely impact segmentation accuracy of deep learning models [8].

Despite these advantages, there are be several challenges involved in generating synthetic alternatives. Training GAN models require ground truth from which to infer new samples, and these models risk overfitting if not provided with sufficient variability. Also, generative models are biologically agnostic which increases the chance that key biological information is lost in the process. Whilst such information can be built in to a computational model, the development of such models may demand disproportionate effort relative to the problem. Computational models follow a rule-based approach for generating an image from reference labels. The use of analytical functions for texturing cell images tend to focus on foreground regions, paying little attention to surrounding regions. However, background features have equal importance in deep learning as models are influenced not only by cell boundaries but also by the surrounding fluorescence context [9].

We propose a sample-based method for generating synthetic cell images suitable for training deep learning models for segmenting 3D fluorescent microscopy data. Our method is efficient, requires little user input, and relies only on low-effort annotations of real images. It addresses two common pitfalls in computational modelling: unnecessary model complexity and choice of fluorescent intensity values. By including biological information in the shape generation process, we present a method that sits between GANs and computational modelling. Our method preserves representative features from a large class of images in fluorescent microscopy: background fluorescence and intensity values across the cell boundary. We show that 3D volume or 2D slice labels from a low number of annotated samples are sufficient to produce training data.

We demonstrate the methods on the Fluo-C3DH-A549 dataset from the Cell Tracking Challenge [10]. This dataset represents a broad class of images for which our method is applicable; cells or cell nuclei where the fluorescent marker is radially distributed. The images from direct application of our method provide useful training data, however we apply several additional steps to refine the synthetic images. A standard UNet trained on our synthetic images achieved excellent segmentation accuracy on real data.

## 2 Methods

### 2.1 Shape Generation

A combination of observations and data from a relevant study was used to simulate labels with similar properties to the cells in the Fluo-C3DH-A549 dataset [11]. The main cell body was initialised in an isotropic domain with size and location to match real data, and a phenotype is selected from wild-type (WT), over-expressing (OE) or phospho-defective (PD). The choice of phenotype determined the number and what type of filopodia were added to the cell body (Fig. 1A).

**Figure 1.**
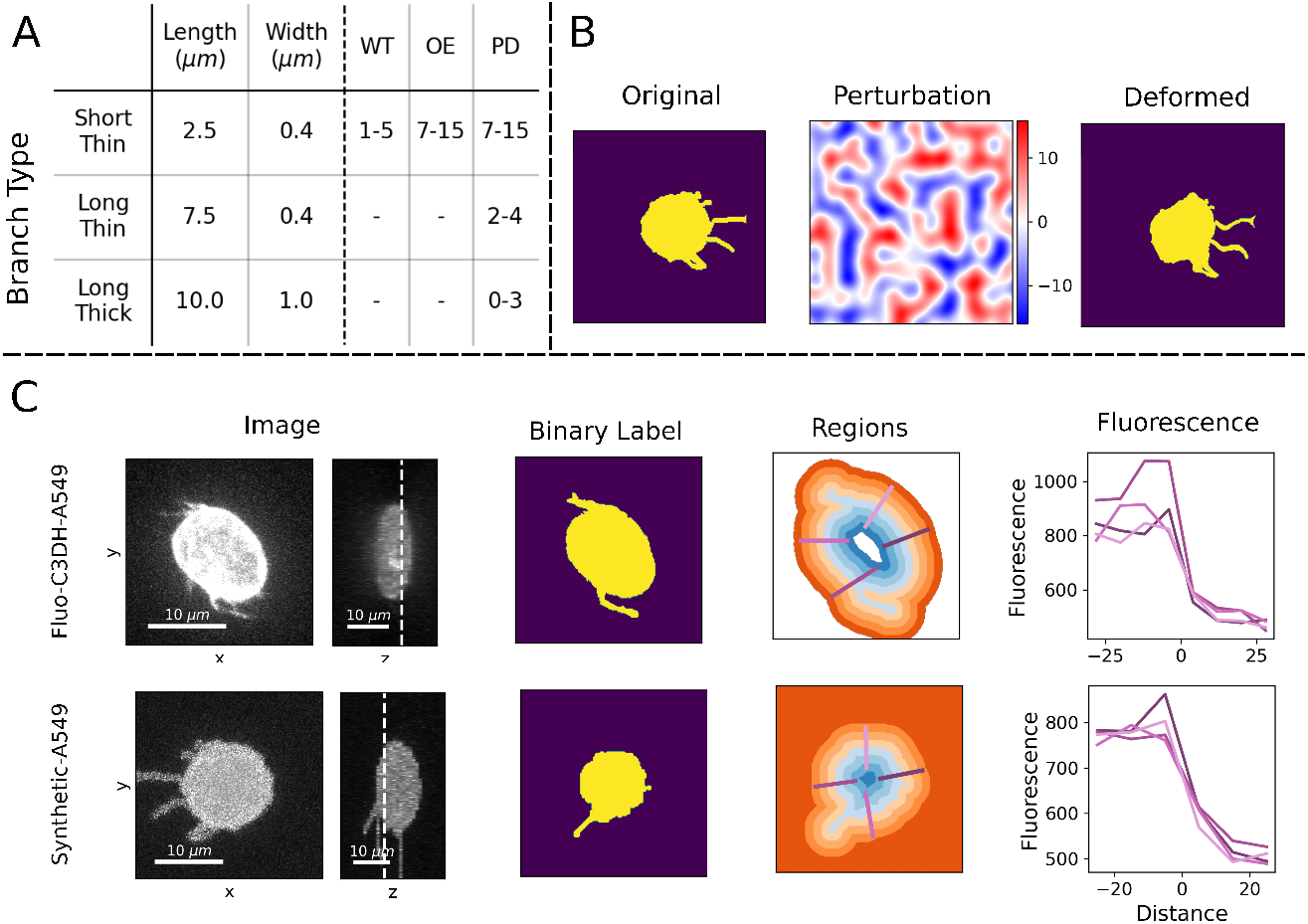
This figure outlines the main steps of synthetic image generation. A) Table of branching filopodia and phenotype parameters. B) A synthetic label example of a PD cell, Perlin noise deformations and deformed version of the example. C) A comparison between real and synthetic images generated by the full method.

When added, a branch was initialised with a direction away from the cell body. Then, depending on the branch type, the branch was built in increments until a target length was reached (see Fig. 1A). At each increment, the direction was perturbed to create a branching effect for the simulated filopodia. Finally, the branch was dilated to a desired width before being added to the final cell label. In the case of a ‘Long Thick’ branch, a tapered effect was implemented such that the base is thicker than at the tip. All simulations were lightweight versions of the computational processes described in [5]. The image label were processed into an anisotropic format to match the data.

Finally, we built a transform for distorting our image labels that uses Perlin noise. The transform follows the same process as the elastic distortions described in [12], replacing the displacements fields with a Perlin noise alternative. This alteration provided greater control when augmenting our synthetic shapes, enabling us to preserve the branch structures (Fig. 1B).

### 2.2 Image Synthesis

We generated synthetic images using the labels as a reference. Instead of assigning pixel intensities with analytical functions, values were sampled from real images. To guide the sampling, we computed distance maps in the 30 annotated real images. Each value in the map corresponded to the signed distance to the label boundary: positive distances outside the boundary and negative distances inside.

Prior to sampling, intensity values from the real images are grouped by regions defined by distance intervals. The intervals were set in increments of *dr* pixels between a minimum (*r*_*min*_) and maximum (*r*_*max*_) distance. Maximum and minimum were imposed to avoid sampling unnecessary regions of the image determined by where the intensity distributions between adjacent regions no longer differed. An example is shown in Fig. 1C, where the regions and corresponding fluorescence profiles are compared along the lines of interest for real data.

Analogous regions were calculated for each synthetic image to determine which region to sample the real intensities from. Pixels at distances greater than *r*_*max*_ were sampled from the furthest background region, while those below *r*_*min*_ were sampled from the most interior region. This sampling regime preserved the fluorescent distribution across the cell boundary which we anticipated as a key feature used in segmentation (see Fig. 1C). In addition, this process yielded more realistic background fluorescence, a feature often neglected in synthetic image generation.

The advantage of this method is that 2D slice annotations can be used in the place of 3D labels. By our sampling routine, a 3D image can be textured using a 2D reference while preserving the key features of the data despite the potential loss of information.

### 2.3 Curvature Weighted Loss

The filopodia are a challenging feature of the image to segment due to the low fluorescence and size compared to the cell body. As a result, a standard cross entropy loss can fail to guide the model to segment the fine protrusions effectively since the reward is too low. To improve the segmentation in these areas, the loss function was weighted at the branches. This approach has been successful in a similar study where the loss function was weighted at the filopodia tips [11]. To localise branches, curvature was estimated at each voxel on the label surface by convolving a sphere of radius 15 pixels. High-curvature regions (H^+^) were defined as all pixels within 15 pixels of a surface voxel where the mask–sphere overlap was below 35%. The union between the image label and regions of high curvature localised the filopodia (see Fig. 2; red regions). The use of curvature was inspired by a curvature-enhanced random walker (CERW) segmentation method which has been successfully applied to this dataset previously [13].

**Figure 2.**
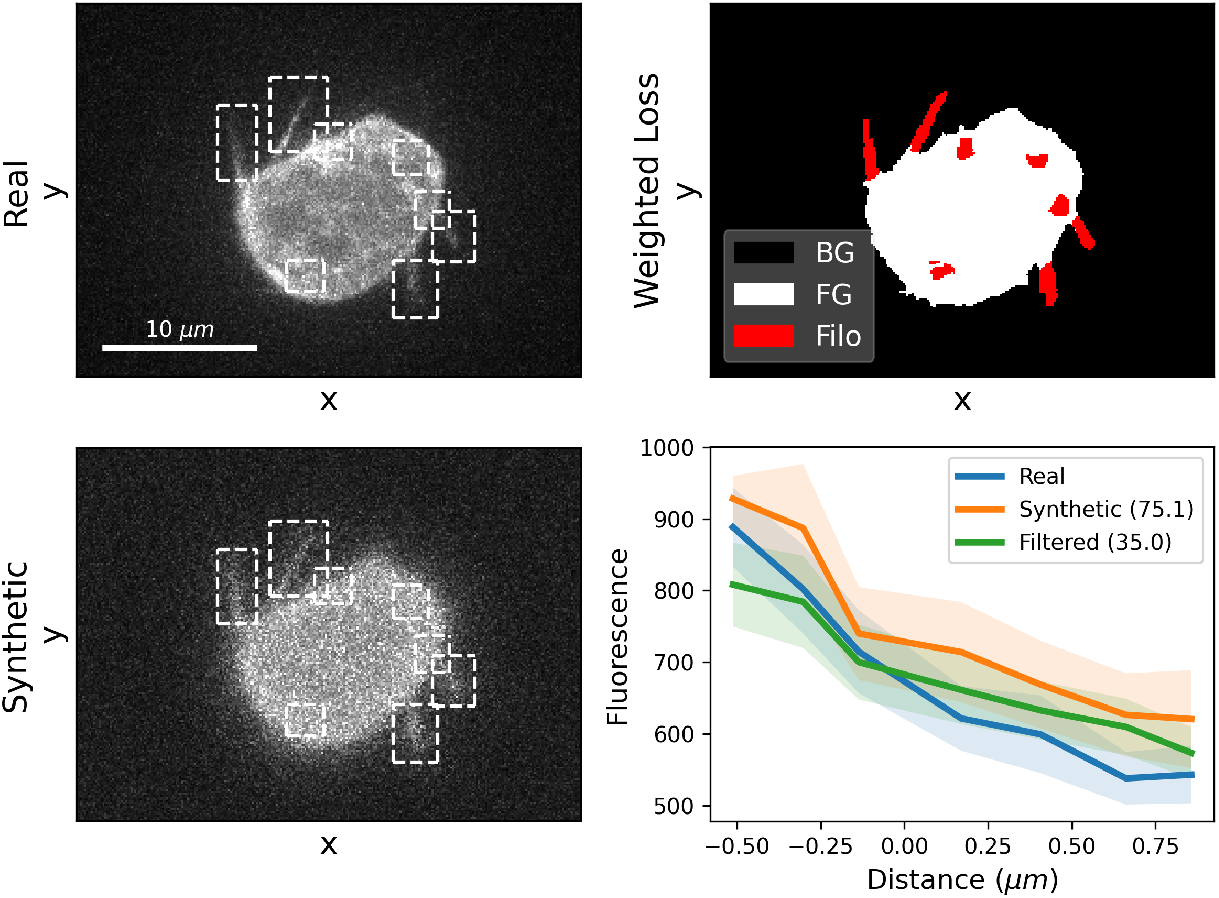
A comparison of filopodia in a real (Top Left) and synthetic image (Bottom Left). The synthetic image is generated by using ground truth annotations with our method. The filopodia are identified through curvature (Top Right). The intensity profile across the filopodia boundary (label surface at 0) are compared for real, synthetic and filtered synthetic images (Bottom Right). Wasserstein distance between the real and synthetic image regions are found in the legend. The lesser value indicates the synthetic image is closer to the real image.

### 2.4 Image Filtering

Applying our sampling method to real image annotations revealed important differences between the real and synthetic images. We plotted the intensity values against the distance to the cell surface around the filopodia, revealing the fluorescence was brighter at the filopodia in the synthetic data compared with the real image (Fig. 2). Therefore, we applied a (3,1,1)-median filter^1^ to the synthetic images to reduce the fluorescence in these regions. The Wasserstein distance between real and synthetic images supported the use of median filtering (see Fig. 2).

Similar analysis revealed a disparity in the fluorescence across the cell boundary below and above the cell body. Slightly higher intensities were found across the cell boundary above the cell in synthetic images compared to the real images. The opposite is observed below the cells which implies a point-spread function. This subtle difference will impact model performance on real data. We rectified this inconsistency by applying an artificial blur in the z-direction with a (2,1,1)-filter. In Fig. 3 we show how a blur parameter, *p* corresponding to a [*p*, 1 − *p*]-filter in the z-direction, was optimised by calculating the Wasserstein distance between the fluorescent distributions in the regions of interest. After evaluating all labelled samples, *p* = 0.3 was chosen.

**Figure 3.**
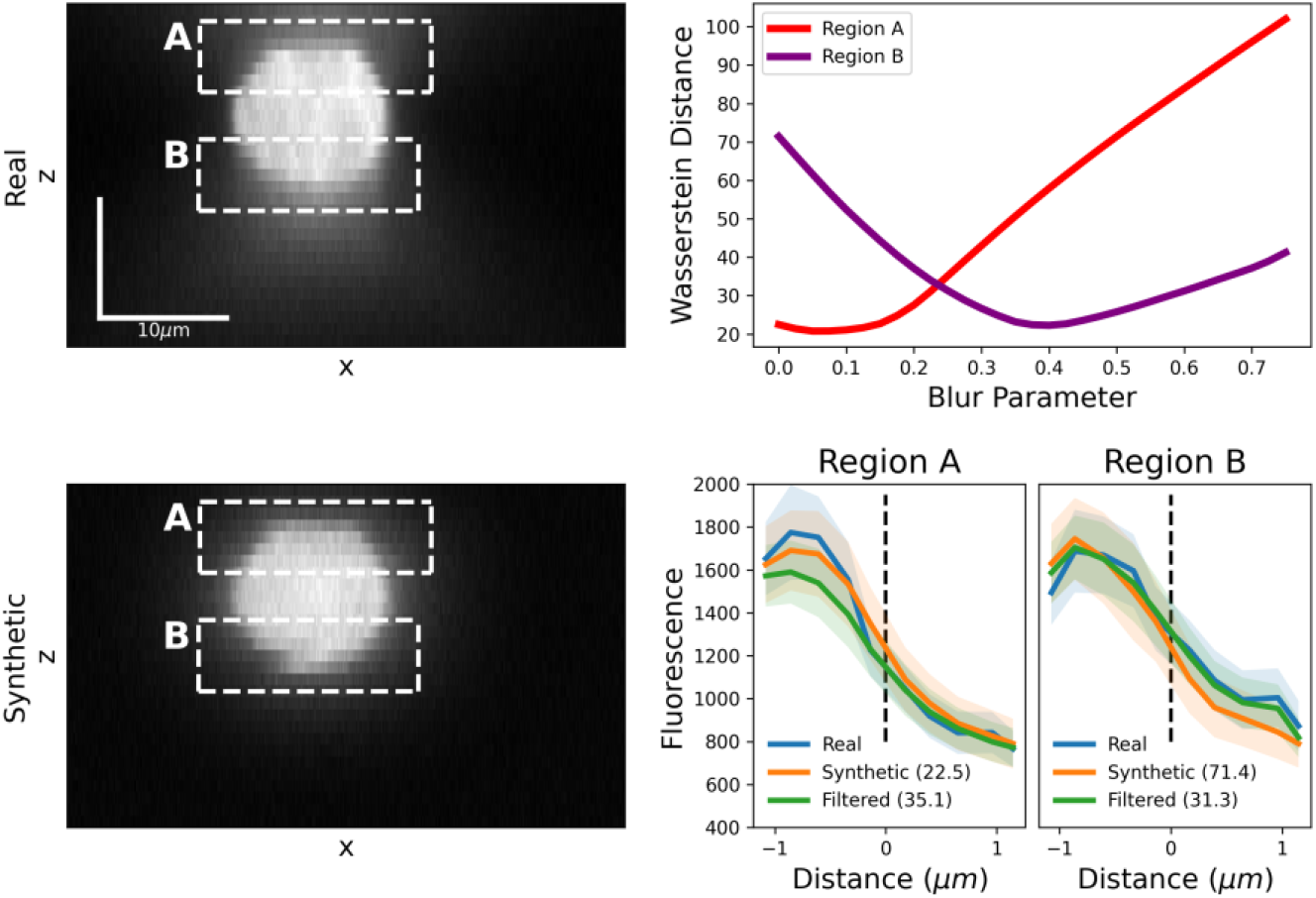
A comparison of fluorescence in real and synthetic images above and below the cell body. The Wasserstein distance is calculated for intensity values in the corresponding regions across real and synthetic images after artificially blurring the latter. An optimal parameter was found to reduce the Wasserstein distance.

### 2.5 Model Training

A standard 5-layer 3D-UNet model was trained on patches of size *D* × *W* × *H* = 16 × 128 × 128, where the first 2 layers use planar convolutions to account for anisotropy. A weighted cross-entropy loss was used, weighting the foreground and filopodia 4-times and 150-times more than the background, respectively. The model was trained for 25,000 epochs before evaluating the segmentation accuracy on the real data every 100 epochs. The accuracy was measured by the mean Jaccard score across all ground truth annotations. During training, a moving average of the accuracy was calculated over 1,000 epochs. The model weights were stored when the moving average Jaccard score peaked during training. Also, we augmented the synthetic images at every training step, applying random Gaussian blur and the elastic deformation described in Section 2.1.

## 3 Results and Discussion

We generated 600 binary shapes following the process outlined in Section 2.1 with equal representation of phenotype. From these reference shapes, two datasets were generated: *VolData* by sampling 3D labels and *SliceData* by sampling 2D slice labels. From each image in the annotated dataset, we selected the xy-slice with the largest labelled region. We generated the two datasets to explore the impact that annotation style has on our method. All models were trained for 100,000 epochs for approximately 48 hours on a single Nvidia A40 GPU. We found that a standard 3D-UNet trained on our synthetic data alone was sufficient to segment real images with an accuracy comparable to the top performing models in the cell segmentation benchmark for this dataset.

### 3.1 Ablation Study

An ablation study was carried out to evaluate the impact of curvature weighted loss and chosen image filtering methods. We evaluated the segmentation accuracy of the trained model by calculating the mean Jaccard score across all annotated real images. Also, using the method described in Section 2.3, we measured the detection and segmentation accuracy at the filopodia (Fig. 2). A detection was considered successful if the model prediction overlapped at least 50% of the region of interest. The CERW was used as a benchmark for our models, as it was the best performing segmentation method on this dataset that did not require training on the labelled images.

Guided by the observations in Fig. 2 and Fig. 3, we trained a complete model on filtered synthetic images using a curvature-weighted loss. Models trained on *VolData* performed excellently on the entire cells, yielding an overall segmentation accuracy of 0.914, as well as detecting 64/133 filopodia with 0.422 segmentation accuracy (Table 1). Training models without the curvature loss function or image filtering resulted in a decrease in all metrics, supporting our choice to include them in the final model. There is a notable decrease in filopodia detection when neither feature is included. All models trained on *VolData* segmented the cells with greater accuracy than the CERW, and a significant improvement was found at filopodia when curvature weighted loss and/or image filtering was used.

**Table 1.**
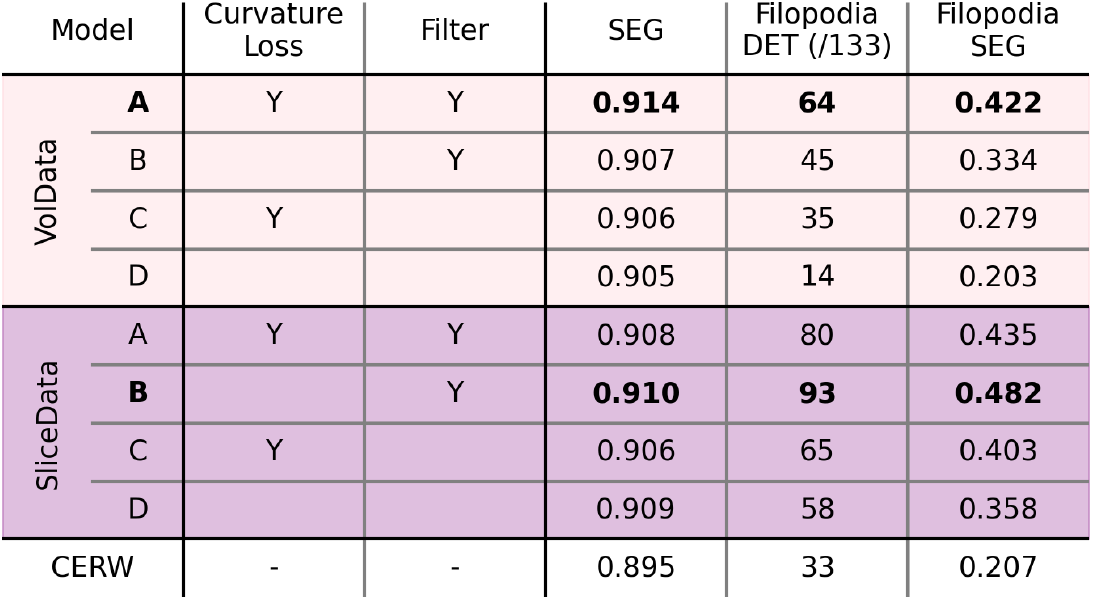
The evaluation metrics for our trained models compared to the CERW. We measure the overall segmentation (SEG) accuracy, detection (DET) accuracy and SEG at filopodia.

### 3.2 Volume vs. Slice Sampling

The complete model trained on *VolData* segmented the overall cells better than the same model trained on *SliceData*. This is evident along the axial direction (See Fig. 4). However, the complete model trained on *SliceData* did a better job at segmenting the filopodia. This result may be due to the axial blur introduced when sampling the 3D annotations. Sampling from slices with less blur results in sharper synthetic images, which may provide better training data for our model.

**Figure 4.**
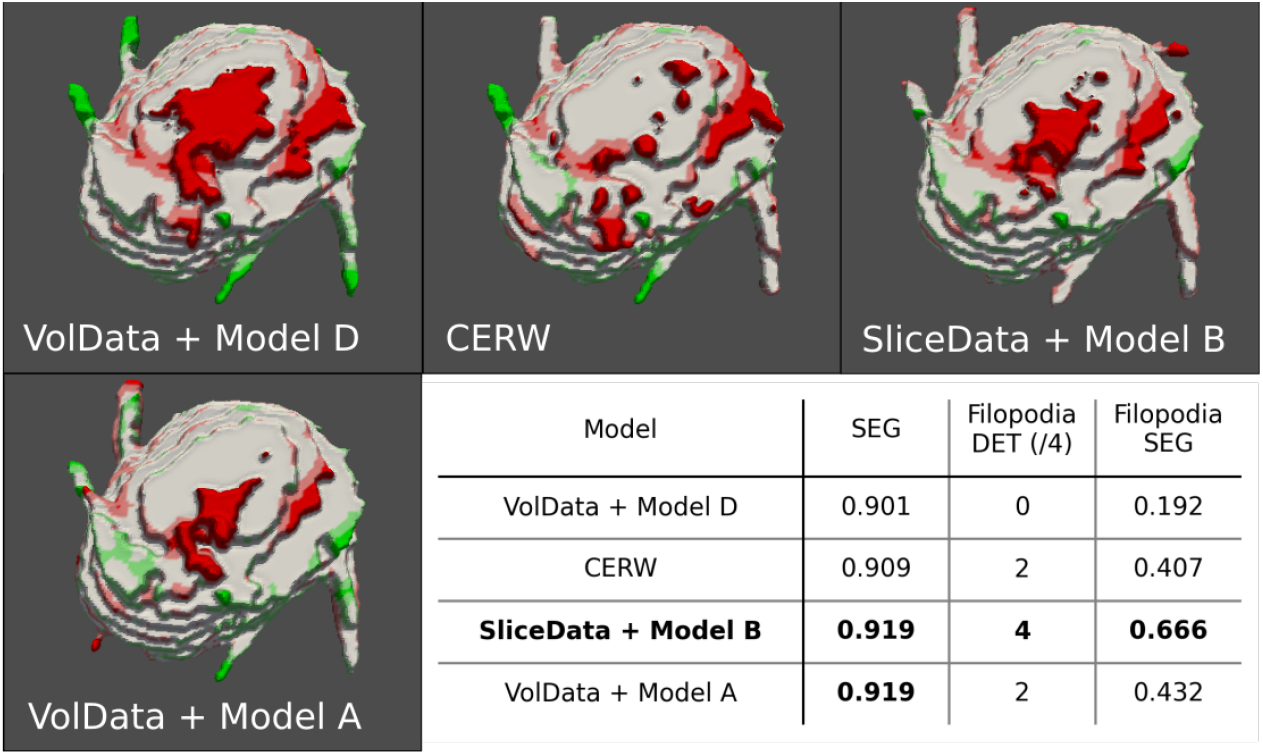
An example of the segmentation results of a real sample for different models (see Table 1). *VolData* + Model D exhibits false positive (FP; red) detection below the cell and false negatives (FN; green) at filopodia. Whilst the CERW improves in these areas, *SliceData* + Model B and *VolData* + Model A achieved the best results.

### 3.3 Cell Tracking Challenge

Finally, we selected the best model trained on *VolData* for the Cell Tracking Challenge submission. The performance on a held-out dataset is evaluated by mean Jaccard score across annotated images and do not consider the filopodia segmentation specifically. Since the ground truth is not available, this featured can not be evaluated for this dataset. Our model achieved the 2nd best segmentation score out of 19 participants (0.904), falling between the CERW score (0.903) and the nnU-Net (0.908) [2].

### 3.4 Discussion

We have demonstrated that segmentation models trained on sample-based synthetic training data compare to top performing models for the Fluo-C3DH-A549 dataset. The models trained on images generated directly from the method performed competitively, which suggested that low-effort slice annotations can be used instead of labourious volume annotations. However, simple filtering techniques informed by ground truth volume annotations improved the results, enabling segmentation accuracies comparable to top performing methods in the segmentation benchmark. Our work showcases the beginnings of a synthetic image generation framework that avoids the complexities involved in alternative approaches without compromising end results.

## 4 Compliance with ethical standards

This is a numerical study for which no ethical approval was required. The dataset used in this study is publicly available on the Cell Tracking Challenge website (celltrackingchallenge.net)

## 5 Acknowledgments

This work was funded through EPSRC/NSF grant EP/X026663/1 to Till Bretschneider.

## Footnotes

1 *z, x, y*−axes matches conventional *D × W × H* model input

